# Attention warps visual-semantic representations across the human cerebellum

**DOI:** 10.64898/2026.10.08.757640

**Authors:** Amanda LeBel, Matteo Visconti di Oleggio Castello, Jack L. Gallant, Richard B. Ivry

**Affiliations:** Department of Neuroscience, University of California Berkeley, Berkeley, CA 94720, USA; Helen Wills Neuroscience Institute, University of California Berkeley, Berkeley, CA 94720, USA; Department of Psychology, University of California, Berkeley, Berkeley, CA 94720, USA

## Abstract

Humans effortlessly shift their attention between objects and events in the environment. These attentional shifts modulate the representation of information in many regions of the cerebral cortex, including those that represent visual-semantic information. Based on recent evidence that the cerebellum also represents semantic information, we hypothesized that attention would also shift semantic representations in the cerebellum. To test this hypothesis, we used fMRI to measure brain activity while participants watched naturalistic movie clips and were instructed to attend to either people or vehicles. We used a voxelwise encoding model framework to recover visual-semantic representations in the cerebral cortex and cerebellum under the two conditions, and quantified the direction and magnitude of visual-semantic tuning shifts between conditions. Attention to different semantic categories had a large effect on visual-semantic tuning in the cerebellum. Representations in some regions shifted toward the attended category, and representations in other regions shifted away from the attended category. Within cerebellar regions that best represent visual-semantic information, the tuning shifts were most pronounced in Crus I and II and in lobule VI. Further, the magnitude of attentional tuning shifts was greater in the cerebellum than in the cerebral cortex, particularly for tuning shifts towards the attended category. These differences suggest that attentional modulation in the cerebellum is different than in the cerebral cortex and may not solely be driven by cerebral inputs.

**Significance Statement:** The cerebellum is increasingly recognized as contributing to cognition, including the representation of meaning. In the cerebral cortex, attention reshapes semantic representations, shifting the tuning of brain activity toward or away from attended categories. Whether the cerebellum shows similar flexibility has been unknown. Using fMRI and voxelwise encoding models, we show that when people search naturalistic movies for humans or vehicles, attention shifts visual-semantic tuning across the cerebellum. These tuning shifts were more widespread in the cerebellum than in the cerebral cortex. Further, the cerebellum had larger shifts towards the attended category than the cerebral cortex. These findings show that cerebellar semantic representations are shaped by attention goals, and suggest that the cerebellum may also contribute to attentional control.

## Introduction

A large corpus of evidence implicates the cerebellum in a broad range of cognitive and affective domains (Schmahmann and Sherman, 1998; Stoodley and Schmahmann, 2010; King et al., 2019; Schmahmann, 2019; LeBel et al., 2021). Within the domain of language, neuroimaging studies have shown that semantic information is represented across both hemispheres of the cerebellar cortex, including crus I/II and lobules VIIB, VIIIA, VIIIB, and IX (Murdoch, 2010; D’Mello et al., 2020; LeBel et al., 2021; Skipper and Lametti, 2021; LeBel and D’Mello, 2023; Casto et al., 2025). These regions have reciprocal connections with association areas of the cerebral cortex (Strick et al., 2009; Caligiore et al., 2017), suggesting that the cerebellum cooperates with the cerebral cortex to represent semantic information in an integrated network, though the specific contributions of these structures remains unclear.

In the cerebral cortex, semantic representations have been recovered using both language comprehension and movie watching tasks (Nishimoto et al., 2011; Huth et al., 2012; Popham et al., 2021). Movie watching tasks have been used to show that the representation of visual-semantic information can shift depending on the state of attention (Kastner et al., 1998; Mazer and Gallant, 2003; David et al., 2008; Çukur et al., 2013; Shahdloo et al., 2020). For example, a voxel that is tuned for inanimate objects when the participant searches for a car may become tuned for people when attention is redirected to search for a person in a crowd. These tuning changes warp the representation of concepts, increasing the cortical representation of attended concepts, while decreasing the representation of unattended concepts. Functionally, these shifts should make it easier to distinguish between attended concepts that are closely related (**Fig 1**). These attention-related shifts in conceptual tuning have also been observed in single neurons (Mazer and Gallant, 2003; David et al., 2008). Moreover, many voxels located within the default mode network show the opposite profile, with their tuning shifting away from the attended target (Çukur et al., 2013). This pattern suggests that attention directed toward specific concepts in the external world increases the representation of those concepts in many brain networks. In contrast, attention decreases the representation of those same concepts in brain networks associated with inner-directed thought.

**Figure 1:**
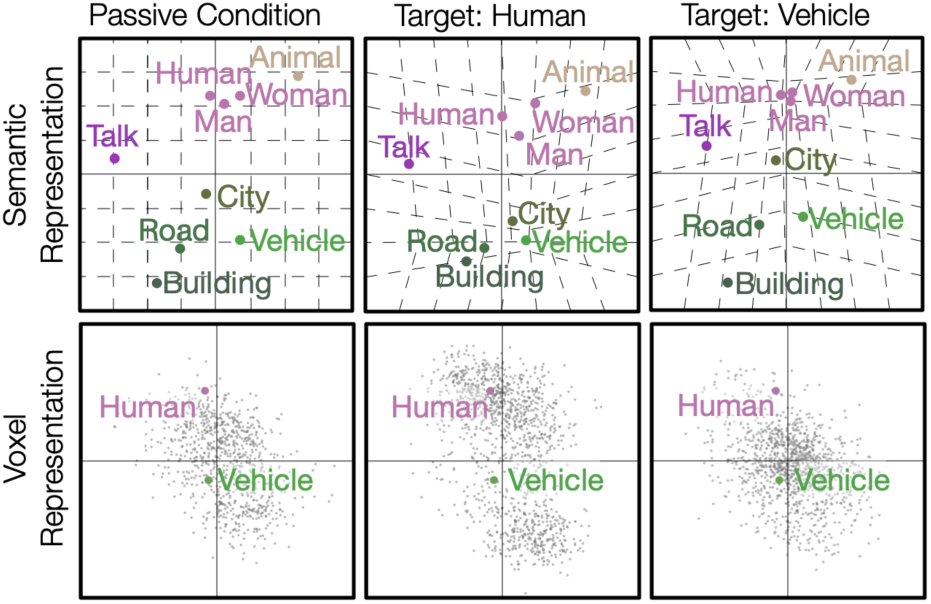
The hypothesized effect of attention on semantic representations. Top: Semantically similar concepts are located nearby one another in semantic space, while dissimilar concepts are located further apart. When attending to a specific category, the semantic space is warped, expanding the area around the attended target. This creates greater separation between semantic topics near the locus of attention, and thus, increasing discriminability of the target (top middle and right panels). Bottom: Warping of semantic space is associated with a shift of the distribution of semantic representations across voxels. Each point in the bottom row depicts the semantic information represented by a single voxel on an idealized two-dimensional semantic space. During passive viewing, the distribution of semantically selective voxels covers the semantic space broadly. When attending to people or vehicles, voxels shift their semantic selectivity toward the attended target, increasing coverage near the attended category.

Here, we ask whether attention expands visual-semantic representations in the cerebellum. Prior studies have shown that attention modulates responses in the cerebellum (Allen et al., 1997; Le et al., 1998), and in subcortical structures such as the superior colliculus and pulvinar (Ignashchenkova et al., 2004; Saalmann et al., 2012). However, these studies have focused on changes in overall activity, showing changes in the cerebellar BOLD response or spiking rate as a function of motor preparatory state (Flierman et al., 2024) or in working memory tasks (Brissenden et al., 2021). No prior study has explored whether and how attention changes the representation of information in the cerebellum.

To examine how attention changes the representation of semantic information in the cerebellum, we adopted the method previously used to study attentional modulation of visual-semantics in the cerebral cortex (Çukur et al., 2013). We collected fMRI data while participants watched a series of movie clips. In separate runs, participants were directed to attend to either the people or the vehicles in the clips. We then used an encoding model framework (**Fig. 2**) (Naselaris et al., 2011; Dupre la Tour et al., 2022; Dupré la Tour et al., 2025) to characterize the visual-semantic representation of each voxel, and to evaluate tuning shifts between the two attention conditions. We used a multi-echo fMRI sequence optimized to increase the signal-to-noise ratio for the cerebellum, and which provides full coverage of both the cerebral cortex and cerebellar cortex. This allowed us to determine whether attentional effects differ between these two brain structures. Furthermore, because past work has all involved language comprehension tasks, the movie watching task provides a novel assessment of semantic representation in the cerebellum.

**Figure 2.**
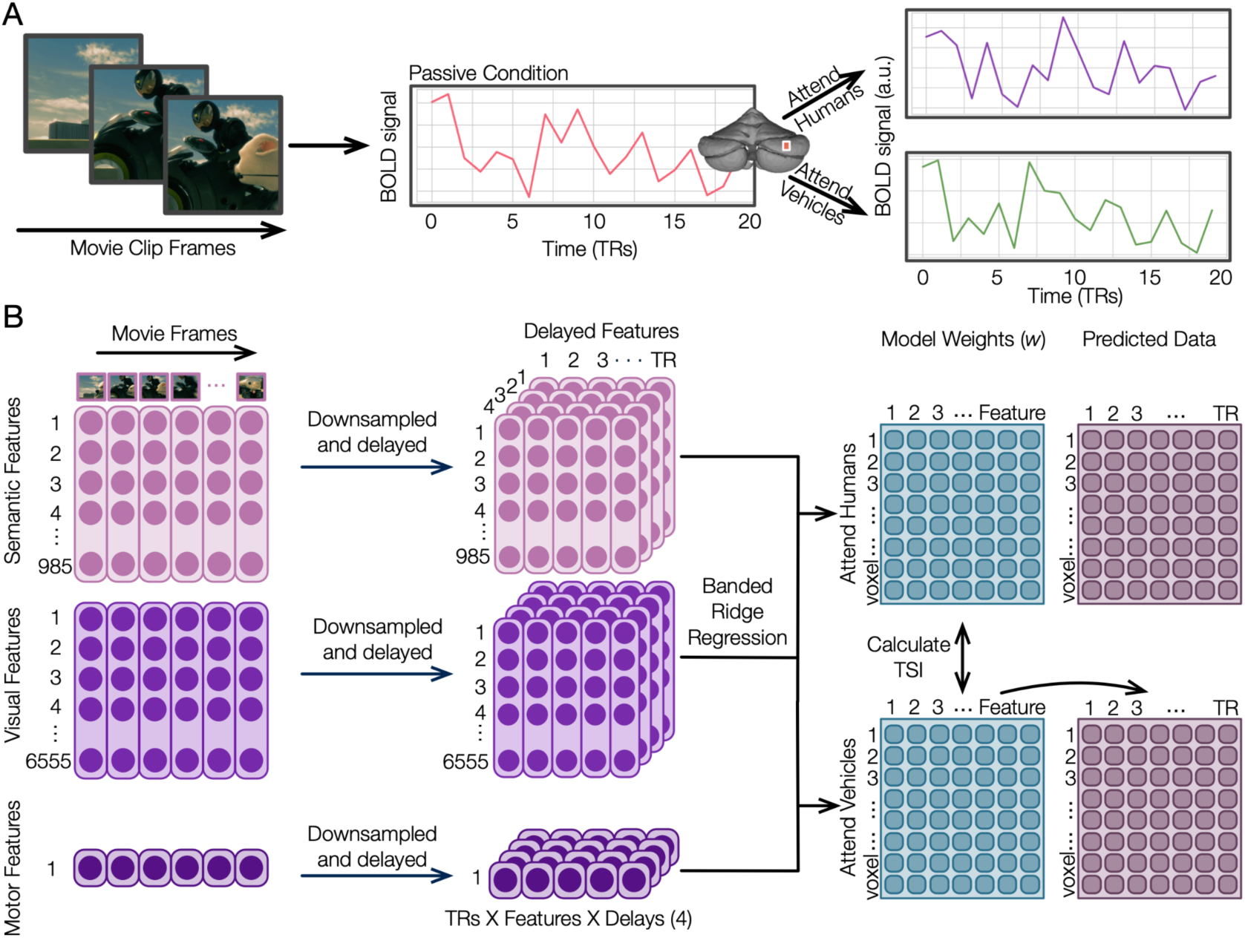
Modeling changes in visual-semantic representation across attention conditions. **A.** To determine how attention changes visual-semantic representations, we estimated the visual-semantic tuning of voxels located in the cerebral cortex and the cerebellum while participants performed a covert visual search task. Participants watched a series of short silent movies while maintaining fixation, and were cued at the beginning of each fMRI run to attend to either “humans” or “vehicles”. **B.** Voxelwise encoding models were constructed in two stages. First, three different types of features were extracted from the video clips. 1) Visual-semantic features that quantify the semantic content in the clips; 2) Motion-energy features that quantify the low-level visual information in the clips; and 3) Motor features reflecting whether or not the participant pressed a button to indicate the presence of a target. In the second stage of the process, banded ridge regression was used to create a separate voxelwise encoding model for each attention condition, and each participant. Voxelwise encoding models were estimated using one training set, and then evaluated using a separate test set. The visual-semantic tuning of each voxel is given by the resulting weight vector for that voxel. The effect of attention on visual-semantic representation was quantified in terms of a tuning shift index across the two attention conditions.

## Methods

### Participants

Functional MRI data were collected from three female participants and one male participant: S01 (female, age 28), S02 (male, age 28), S03 (female, age 25), S04 (female, age 27). All participants were healthy and had normal or corrected to normal vision, they were highly trained, motivated, and experienced in fMRI, and they were aware of the experimental manipulations. The experimental protocol was approved by the Institutional Review Board at the University of California, Berkeley. Written informed consent was obtained from all participants.

### Experimental overview

Each participant took part in a total of four scanning sessions. Functional localizers and anatomical data were collected in the initial two sessions. Functional scans for the main experiment were collected in the remaining two sessions. The experimental stimulus and task used here were identical to those described in Cukur et al, 2013. However, all of the fMRI data used in this report were newly collected for this project.

### Stimulus

The stimulus set was primarily obtained from Hollywood movie trailers that were cut to create a series of shortclips (10-20 s) spanning a broad range of settings and events. These short clips were spliced together to create three unique movies, each lasting 600 s. The combined 1800 s of movies included 450 s of frames containing only humans or only vehicles, 450 s of frames containing both humans and vehicles, and 450 s of frames containing neither humans or vehicles. A fixation point was superimposed on the movies and its color was alternated at 1Hz to maximize visibility. The stimuli were presented at a rate of 15 Hz using an MR safe projector (Avotec). At the beginning of each scanning run, participants were cued to attend to one of two potential targets (“humans” or “vehicles”). The attentional targets appeared in many settings, positions, sizes, and viewpoints. The same three movies were used in both attention conditions.

### Task

The two scanning sessions of the main experiment were each composed of six 10-min runs. At the start of each run, participants were instructed to fixate at a center fixation point, and were cued to covertly search for either “humans” or “vehicles” while watching movie clips. To ensure participants maintained vigilance, they were instructed to press a button every time a target from the designated category appeared on the screen, and to keep the button pressed for as long as the target remained on the screen. All participants used their right hand to respond. This task was repeated for three “attend human” runs and three “attend vehicle” runs per session. To avoid sampling bias in the stimulus, the same three movie clips were presented in both attention conditions and the conditions were alternated within each session. The three movie clips were first shown with the conditions “vehicle”, “human”, “vehicle”, and then the three movie clips were repeated in the same order with the conditions “human”, “vehicle”, “human”. Within each run, the presence of each target was balanced and randomized. One quarter of the clips contained only humans, one quarter of the frames contained only vehicles, one quarter of the frames contained both humans and vehicles, and one quarter of the frames contained neither humans or vehicles. To increase statistical power, the same procedure was repeated in a second scanning session.

### MRI data collection

MRI data was collected on a 3T Siemens Trio scanner located in the UC Berkeley Brain Imaging Center. For the two main sessions, a multi-echo gradient-echo EPI sequence and 32 channel head-coil were used to collect functional data. This sequence had the following parameters: repetition time (TR) = 1.50 s, echo times (TE) = [14.6 ms, 34.47 ms, 54.34 ms], flip angle = 71°, multi-band factor (simultaneous multi-slice) = 3, PAT = 2, voxel size = 2.6mm x 2.6mm x 2.6mm (slice thickness = 2.6mm), matrix size = (98, 98), field of view = 257 mm, anterior-posterior phase-encoding direction (Visconti di Oleggio Castello et al., 2021). The field of view encompassed the entire cortex and the cerebellum in all participants. Before every functional scan, two spin-echo volumes with opposite phase-encoding direction were acquired to compute fieldmap estimates.

The two localizer sessions for each participant were previously collected using a different pulse sequence as part of other experiments. The localizer sequence had the following parameters: repetition time (TR) = 2 s, echo time (TE) = 34 ms, flip angle 74°, voxel size = 2.24 x 2.24 x 3.5 mm^3^, field of view = 224 x 224 mm^2^, and 32 axial slices to cover the entire cerebrum. This scanning sequence did not have full coverage of the cerebellum, so functional ROIs in the cerebellum were defined at the group level from a previous dataset (see below) (LeBel et al., 2023). Using a different sequence for the localizer tasks does not affect the main results reported here because the functional ROIs are used solely for visualization purposes and are not used in any statistical analysis. Four anatomical scans were collected for each participant following the FreeSurfer morphometry protocol. The sequence was a T1-weighted multi-echo MP-RAGE with voxel size = 1mm x 1mm x 1mm, run on the same scanner as was used to acquire localizers.

### fMRI preprocessing

Each functional run was motion-corrected, distortion-corrected, and aligned to the participant’s T1-weighted volume. All spatial transformations were estimated on the volumes acquired at the first echo, which provides higher tissue contrast. The same spatial transformations were then applied to the volumes acquired at the other two echoes. For each run, motion-correction parameters were estimated with MCFLIRT from FSL 6.0 (Jenkinson et al., 2002; Woolrich et al., 2009). The first-echo, single-band reference volume was used as a reference for motion correction. FSL’s TOPUP (Andersson et al., 2003) was used to estimate the distortion correction transformation from the two spin-echo volumes acquired with opposite phase encoding directions. The distortion-correction transformation was then applied to the single-band reference volume to produce an unwarped reference volume. The bbregister function from Freesurfer was used to align the unwarped reference volume to the participant’s high-resolution T1-weighted volume (Greve and Fischl, 2009). ANTs (Avants et al., 2011) with Lanczos interpolation was used to apply the motion-correction, distortion-correction, and alignment transformations in a single step. For each run, this process produced a set of motion-corrected, distortion-corrected, and anatomically aligned volumes from three echo times. Finally, the @compute_OC_weights from AFNI was used to optimally combine these three volumes into a single volume. The resulting spatially preprocessed data were then temporally detrended and denoised with nuisance regressors estimated from aCompCor (Behzadi et al., 2007). The anatomical mask for aCompCor included white matter voxels from both the cerebrum and cerebellum. After denoising, scipy.signal.resample was used to temporally resample the data in each voxel from a 1.5 s to a 2 s TR. Finally, the data were temporally z-scored. All preprocessing pipelines were implemented in Python using nipype (Gorgolewski et al., 2011, 2016), sdcflows (https://github.com/nipreps/sdcflows), niworkflows (https://github.com/nipreps/niworkflows), and custom code.

### Cortical surface reconstruction and visualization

Freesurfer 6.0 (Dale et al., 1999) was used to generate high-resolution cortical surfaces from the T1-weighted anatomical scans. Before surface reconstruction, anatomical surface segmentations were manually inspected and corrected. Blender was used to remove the medial wall and make relaxation cuts for flattening.

### Cerebellar surface reconstruction and visualization

The SUIT toolbox (Diedrichsen, 2006) was used to isolate the cerebellum from the rest of the brain in the T1-weighted anatomical image. The isolation mask was manually inspected and corrected. The SUIT registration algorithm was used to spatially transform the participant-specific anatomical maps of the cerebellum into the template SUIT space. The resulting transformation was used to map the voxelwise encoding results to the SUIT space only for visualization. All analyses were performed in the participant’s native functional space.

To visualize imaging results on both the cortical and cerebellar surfaces, the line_nearest projection scheme in pycortex (Gao et al., 2015) was used to project the values in each voxel onto the surface. This projection samples the functional data at 32 evenly spaced intervals between the inner (white matter) and outer (pial) surfaces. Samples are taken according to nearest-neighbor interpolation, and each sample is given the value of its enclosing voxel. The resulting samples are then averaged and assigned to the vertex.

### Regions of Interest and Functional Localizers

To provide functional landmarks in both the cerebrum and cerebellum, separate localizers were used to identify known regions of interest (ROIs) in each participant. Three different tasks were used to define the ROIs: a visual category localizer, a motor localizer, and a retinotopic localizer. Cerebral ROIs were identified in individual participants. Cerebellar ROIs were identified at the group level to compensate for the partial coverage of the scanning sequence that was used for the localizer tasks. Note that in this study neither the cerebellar ROIs nor cortical ROIs are used for specific analyses; they are only used to aid in the identification of functionally defined anatomical landmarks.

### Visual category localizer

Data were collected in six 4.5 minute scans consisting of 16 blocks of 16 seconds each. During each block 20 images of either places, faces, bodies, household objects, or spatially scrambled objects were displayed. Participants were asked to detect when the same image was presented twice in a row. This localizer was used to define ROIs in the cerebral cortex corresponding to the fusiform face area (FFA) (Kanwisher et al., 1997), occipital face area (OFA)(Kanwisher et al., 1997), extrastriate body area (EBA) (Downing et al., 2001), parahippocampal place area (PPA) (Epstein and Kanwisher, 1998), and the occipital place area (OPA).

### Motor localizer

Data was collected during 2 identical 10-minute scans. The participant was cued to perform six different tasks in a random order in 20-second blocks. The cues were the words ‘hand’, ‘foot’, ‘mouth’, ‘speak’, or ‘rest’ presented at the center of the screen, except for the saccade cue which was presented as a random array of dots scattered across the display. For the hand cue, participants were instructed to make small finger-drumming movements for the 20 s cued period. For the foot cue, the participants were instructed to make small foot and toe movements. For the mouth cue, participants were instructed to make small vocalizations of nonsense syllables such as *balabalabala*. For the speak cue, participants were instructed to self-generate a narrative without vocalization. For the saccade cue, participants were instructed to move their eyes around the dot display. The motor localizer was used to define ROIs in both the cerebral cortex and the cerebellum. The cortical ROIs included primary motor and somatosensory areas for the hands, feet, and mouth; supplemental motor areas for the hands and feet, secondary motor areas for the hands, feet, and mouth, the ventral premotor hand area; frontal eye field and intraparietal sulcus visual areas; Broca’s area and the superior ventral premotor area (sPMV) speech area. The cerebellar ROIs were computed by resampling the motor localizer model weights to SUIT space, and then averaging the models across participants. Motor areas for the hand, mouth, foot, and saccade tasks were defined in the posterior and anterior lobe as functional landmarks.

### Retinotopic localizer

Data was collected following the protocol detailed in a previous publication (Hansen et al., 2007). In brief, during the retinotopic localizer two distinct visual stimuli were used, a wedge stimulus and a ring stimulus. The wedge stimuli consisted of eight adjacent 45° wedges and the ring stimuli consisted of four concentric rings. Both wedges and rings were filled with overlapping polar and hyperbolic non-cartesian gratings. Gratings were placed randomly within wedges or rings. The spatial phase of each grating was advanced slightly on each image frame to create dynamic stimuli. Spiral gratings rotated clockwise or counterclockwise, and hyperbolic gratings expanded or contracted. The color, spatial frequency, and apparent velocity of each grating were assigned randomly. During a functional run, wedges or rings were turned on and off to map foveal angle and foveal eccentricity, and to map early visual cerebral cortical regions of interest including V1, V2, V3, V3A, V3B, V4, hMT+, LO, and V7.

### Experimental Design And Statistical Analyses

#### Feature spaces

Three feature spaces were extracted from the stimuli and were used to model brain responses: a semantic feature space (*eng1000*), a low-level visual feature space (*motion energy*), and a motor feature space.

To generate the semantic feature space, each 1 s clip of the movies was manually labeled using terms from the WordNet lexicon.(Fellbaum and Miller, 2021) Three naive raters performed the labeling, and potential conflicts were resolved by conferral among the raters. These WordNet labels were then converted to a word co-occurrence semantic model (*eng1000*). We opted to perform this conversion to allow for comparison between this work and previous semantic models of the cortex and cerebellum (Huth et al., 2016; LeBel et al., 2021, 2023). To perform the conversion from WordNet to *eng1000*, each label within each 1 s clip was assigned to the corresponding vector from the *eng1000* space. The resulting vectors for all labels in each 1 s clip were averaged together, and a 3-lobe Lanczos filter was used to downsample the vectors to the fMRI sampling frequency.

To generate the low-level visual feature space, each frame of the movie was passed through a visual motion-energy model (Adelson and Bergen, 1985; Watson and Ahumada, 1985; Nishimoto et al., 2011). This motion-energy model consists of 8395 space-time quadrature Gabor filter pairs. The filters were selected to cover the entire image space, and reflected a wide range of preferred receptive field sizes, orientations, and spatiotemporal frequencies. The spatial frequencies were 0, 2, 4, 8, 16, and 32 cycles per degree. The orientations were 0, 45, 90, 135, 180, 225, 270, and 315 degrees. The temporal frequencies were 0, 2 and 4 Hz (Huth et al., 2016; LeBel et al., 2023). A 3-lobe Lanczos filter was used to downsample the resulting outputs of all filters to the fMRI sampling frequency.

To generate a motor feature space, each participant’s button presses were converted to a binary vector. The button presses were recorded at a rate of 15 Hz and were decimated to the fMRI sampling frequency. This resulted in a binary one-dimensional vector consisting of ones at the time points when the button was pressed and zeros otherwise.

#### Fitting voxelwise encoding models

Two separate voxelwise encoding models were created for each participant, one for the “attend humans” condition and one for the “attend vehicles” condition. To fit these models, the stimulus matrix for each of the three feature spaces was z-scored separately within each run, and then concatenated across runs. To account for the delay in the hemodynamic response function (HRF), four delayed copies of the stimulus matrix (using delays of 1, 2, 3, and 4 time points) were concatenated together using a finite impulse response procedure. To account for possible stimulus correlations between the three feature spaces, banded ridge regression (Dupré la Tour et al., 2022) was used to predict the BOLD data from the delayed feature matrix. Banded ridge regression is a variant of ridge regression in which each feature space is regularized by a separate hyperparameter. By regularizing each feature space separately, banded ridge regression can account for correlated feature spaces and improves model prediction accuracy (Nunez-Elizalde et al., 2019; Dupré la Tour et al., 2022).

Voxelwise encoding models were optimized, fit, and evaluated in separate train, validation, and test sets. Nested cross-validation was used to generate these datasets (See **Supplementary Figure S.1)**. First, the entire dataset was split into a training set and a test set (outer cross-validation fold). Then, the training set was further divided into a smaller training set and a validation set (inner cross-validation fold). The smaller training set was used to fit voxelwise encoding models with a range of regularization hyperparameters. Hyperparameters were selected from 20 log-spaced values between 10 and 10^20^ according to random search with 400 iterations. Each of these 400 models was evaluated in the validation set. The process was repeated for all possible validation sets. Then, the hyperparameters yielding the highest average prediction accuracy (R^2^) in the validation set were selected, separately for each voxel. Finally, a voxelwise encoding model was fit on the full training set in the outer cross-validation fold, and evaluated on the test set. The entire procedure was repeated for all possible combinations of training and test sets, and model prediction accuracy was averaged across test sets.

Because each unique movie was presented twice for each attention condition (once in each fMRI scanning session), the nested cross-validation procedure was performed by holding out the two repeats of a single movie stimulus as a test set. Therefore, the training set in the outer cross-validation fold included data from four scanning runs (corresponding to two unique movie stimuli), and the test set included data from two scanning runs (corresponding to one unique movie stimulus). The same scheme was used for the training/validation splits. This scheme ensures that the training/validation/test splits are independent in terms of both collected data and presented stimuli. This nested cross-validation procedure is illustrated in **Figure S.1.**

Because an independent regularization hyperparameter was used for each voxel, the scale of the model weights may differ between voxels. This difference in scale can complicate comparison of model weights across voxels with different regularization hyperparameters. To mitigate these differences in scale, the norm of the model weights was rescaled according to the average validation-set R^2^ score within the training set (Meschke et al., 2023). This rescaling step effectively ensures that the scale of the model weights is comparable across voxels. This step was performed independently for each cross-validation split before averaging the model weights for further analysis.

Model prediction accuracy was quantified with the correlation coefficient (r) by comparing the model prediction, *Y^^^___*, to the held out test data, *Y*. (Similar results were obtained by using the coefficient of determination R^2^.) Because the voxelwise encoding models are linear, the model prediction can be decomposed into the split prediction of each separate feature space, such that:

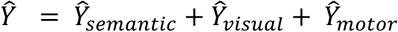

To evaluate the contribution of each feature space to the model prediction, the overall model prediction accuracy was decomposed with the split correlation coefficient (r_split) (St-Yves and Naselaris, 2018; Dupré la Tour et al., 2022). The split correlation coefficient ensures that:

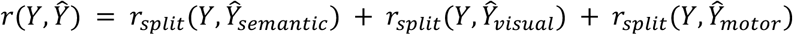

#### Statistical significance of model prediction accuracy

Statistical significance of model prediction accuracy was evaluated with permutation tests. Because a nested cross-validation procedure was used to estimate and evaluate voxelwise encoding models, a separate null distribution was generated for each of the held-out test sets. For each test set, the time series for each voxel was randomly shuffled in blocks of 10 TRs and the correlation with the predicted time series was recalculated. This was done for 1000 permutations of each held-out test set. Finally, the resulting null distributions were pooled to compute empirical p-values in each voxel. These p-values were FDR corrected to account for multiple comparisons (Benjamini and Hochberg, 1995). A threshold of *q*(*FDR*) < 0.05 was used as a criterion to define significantly well predicted voxels.

#### Characterization of tuning shifts

The primary aims of this study were to determine if semantic tuning in the cerebellum is modulated by attention, and to evaluate how any such tuning shifts compare with those observed in the cerebral cortex. To determine whether modulation occurs, it is necessary to quantify shifts in semantic tuning across attention conditions (**Figure 1**). Following previous work (Çukur et al., 2013), the extent of tuning shifts was measured by fitting voxelwise encoding models to the BOLD time series while participants were instructed to attend to either “humans” or “vehicles”. The resulting model weights measure the tuning of each voxel towards specific visual-semantic features. To quantify the amount of semantic tuning specifically towards humans and vehicles, we first created a template weight vector for the “humans” semantic category and a template weight vector for the “vehicles” semantic category. The “humans” template vector was created by averaging the vectors corresponding to the words “person”, “man”, “woman”, “child”, “athlete”. The “vehicles” template vector was created by averaging the vectors corresponding to the words “ship”, “airplane”, “truck”, “car”, “vehicle”. These template vectors were then correlated with the semantic model weights in each voxel to quantify the tuning strength towards vehicles and humans.

Mathematically, in each voxel the tuning strengths are defined as

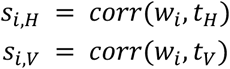

Here, *s_i_*_,*H*_ is the tuning strength for humans and *s_i_*_,*V*_ is the tuning strength for vehicles during each attention condition *i* (*i* = *H*: search for humans, and *i* = *V*: search for vehicles). *w_i_* is the voxelwise weight vector during condition *i*, and *t_H_* and *t_V_* are the templates for human and vehicles respectively.

To summarize the magnitude of attentional shift occurring in each voxel, the measured tuning strengths were used to compute a tuning shift index (TSI) (Çukur et al., 2013). The TSI is computed as:

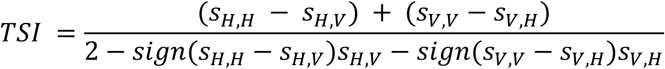

The numerator measures the difference in tuning strength for the attended versus unattended category, summed across the two attention conditions. The denominator scales the TSI to [-1, 1]. If the voxel tuning does not change between the two attention conditions, the TSI is 0. If the voxel tuning shifts towards the attended target, the TSI is positive. If the voxel tuning perfectly matches the attended target template in both attention conditions, the TSI is 1. In contrast, if a voxel tuning shifts away from the attended target, the TSI is negative. If a voxel tuning shifts towards the unattended target in both attention conditions, the TSI is −1.

#### Statistical significance of tuning shifts

Statistical significance of tuning shifts was computed with permutation tests. To generate a null distribution for each voxel, the model weights were randomly shuffled 1,000 times separately for the “attend human” and “attend vehicle” conditions. The TSI was then calculated for each permutation, resulting in a null distribution of TSI values for each voxel. The null distribution was used to calculate an empirical p-value for each voxel, and the p-values were FDR-corrected to account for multiple comparisons (Benjamini and Hochberg, 1995). A threshold of *q*(*FDR*) < 0.05 was used as a criterion to define voxels with a significant tuning shift.

#### Evaluation of tuning shifts within cerebellar ROIs

To compare tuning shifts across anatomical regions in the cerebellum, tuning-shift analyses were also performed within participant-specific anatomical ROIs. Lobular ROIs within each participant were defined by means of an automatic parcellation algorithm (ACAPULCO (Han et al., 2020)). To compute an average TSI value for each lobular ROI, voxelwise TSI values were averaged only for voxels with significant tuning shifts (q(FDR) < 0.05). To ensure that the average TSI value was not biased by voxels with low model prediction accuracy, before averaging the TSI value of each voxel was weighted by its cross-validated visual-semantic split prediction accuracy (R^2^) in the training set. The same procedure was also used to compare TSI values between the whole cerebellum and the whole cortex.

## Results

The goals of this study were to examine whether visual-semantic representations in the cerebellum are modulated by attention, and to compare any such shifts to those observed in the cerebral cortex. We collected BOLD data while participants were cued to covertly attend to either the “humans” or “vehicles” in a series of silent naturalistic short clips (60 minutes of data for each attention condition). To quantify attentional tuning shifts, we fit separate voxelwise encoding models for each of the two attention conditions. Each voxelwise encoding model was based on a visual-semantic feature space that was shown previously to capture both linguistic and non-linguistic semantic representations in the cortex and in the cerebellum (Huth et al., 2012, 2016; LeBel et al., 2021, 2023; Popham et al., 2021). Motor and visual motion energy feature spaces were also included in the encoding models to minimize potential confounds arising from motor responses or by low-level visual information in the stimuli.

To evaluate model prediction accuracy, a held-out data set was used to calculate the correlation (r) between the obtained functional responses and those predicted by the fit voxelwise encoding models (see Methods, **Figure 2**). **Figure 3A** shows the model prediction accuracy for the “attend human” condition in both the cerebral cortex and in the cerebellar cortex. (Overall model prediction accuracy was similar between the two attention conditions, see **Supplementary Figure S.2** for “attend vehicle” condition.) Voxels throughout the visual cortex, along the temporal lobe, and in regions of the frontal lobe are significantly predicted (q(FDR) < 0.05) by the encoding model. In the cerebellum, voxels in crus I/II bilaterally, along the vermis, as well as voxels in lobules VI, VIIB, and IX are significantly predicted (q(FDR) < 0.05) by the model. These results indicate that the encoding models accurately capture representations within the cerebral cortex and cerebellar regions, consistent with previous studies (Çukur et al., 2013; van Es et al., 2019; LeBel et al., 2021). Because our goal was to compare functional representations across attention conditions, all subsequent analyses focused only on voxels that were significantly predicted in both attention conditions.

**Figure 3.**
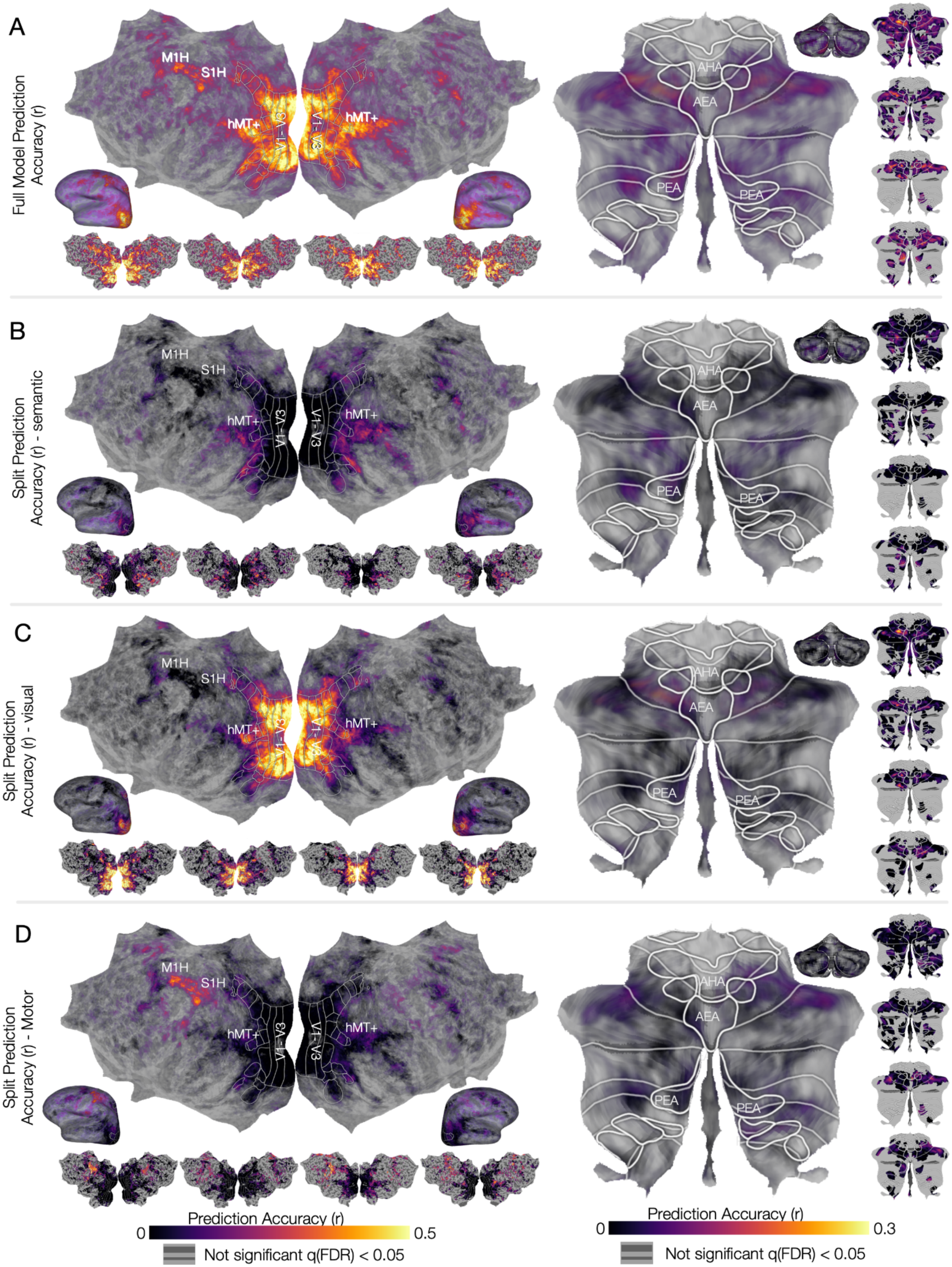
Prediction accuracy of voxelwise encoding models in the cerebral cortex and the cerebellum in the “attend human” condition. To identify regions of the cerebral cortex and the cerebellum that selectively represent semantic information, banded ridge regression was used to fit voxelwise encoding models with three feature spaces: visual-semantic, low-level visual, and motor. A nested cross-validation procedure was used to train and test models in separate data sets. To identify cerebral and cerebellar areas that are well predicted by the voxelwise encoding model, model prediction accuracy (correlation, *r*) was averaged across participants, and projected onto the fsaverage and SUIT surfaces respectively. **A.** Model prediction accuracy (correlation, *r*) for the full model, comprising all three feature spaces, is plotted on flattened cortical and cerebellar surfaces for the group (center, large) and in each individual participant (small). The full model in the cerebral cortex shows high prediction accuracy through the visual cortex, along the temporal lobe, and primary somatomotor cortex. Within the cerebellum, the full model shows high prediction accuracy in lobules VI, Crus I/II, and lobule VIIB. To disentangle the contribution of each feature space to model predictions, the split prediction accuracy was plotted on the same cerebral and cerebellar surfaces (B, C, and D). The split prediction accuracy can be interpreted as the variance explained by each feature space. **B.** Split prediction accuracy maps for the semantic feature space. The semantic features best explain variance in the temporal and frontal lobes in cerebral cortex, and in Crus I/II and lobule VIIB in the cerebellum. **C.** Split prediction accuracy maps for the low-level visual feature space. The low-level visual feature space best explains variance in the visual cortex, and in lobules VI and vermis in the cerebellum. **D.** Split prediction accuracy maps for the motor feature space. The motor feature space best explains variance in the left primary somatomotor cortex, and in right lobules I-IV and VI in the cerebellum. These results indicate that separate regions in the cortex and cerebellum represent semantic, low-level visual, and motor information during the naturalistic visual search task. Representations in voxels that are well predicted by the semantic feature space can be reliably interpreted to determine changes in semantic tuning across attentional states.

To identify voxels that are well predicted by the visual-semantic feature space, the split model prediction accuracy was used to determine the contribution of each individual feature space to model prediction. Split prediction accuracy captures the relative amount of variance captured by each feature space for each voxel. **Figure 3B** shows the split prediction accuracy for the visual-semantic features in the cerebral cortex and cerebellum. Within the cerebral cortex, the semantic feature space explains variance primarily in the temporal lobe, parietal lobe, and portions of the frontal lobe. In the cerebellum, the semantic feature space explains variance primarily in crus I and II, and in lobule VI. These results suggest that regions in both the cerebral cortex and cerebellum represent semantic information, consistent with previous findings (Huth et al., 2012, 2016; de Heer et al., 2017; LeBel et al., 2021).

**Figure 3C** shows the split prediction accuracy for the low-level visual feature space in the cerebral cortex and cerebellum. Within the cortex, the low-level visual feature space explains variance primarily in V1-V3, in extrastriate visual areas such as hMT+, and in the frontal eye-movement areas such as FEF and SEF. Within the cerebellum, the low-level visual feature spaces explains variance primarily in lobules VI, VIIB, and along the vermis. These cerebellar regions are adjacent to known eye motor regions, the anterior eye area (AEA) and posterior eye area (PEA). **Figure 3D** shows the split prediction accuracy for the motor feature space in the cerebral cortex and cerebellum. In the cortex the motor feature space explains variance largely in the left primary somatosensory and motor hand region (S1H and M1H respectively), as well as in secondary somatomotor regions in the insula. Within the cerebellum, the motor feature space explains variance within the right lobules V and VI. Note that all participants responded with their right hand during the task, which explains the lateralization in the cortex and cerebellum.

Taken together across all feature spaces, these results show that voxels in regions of both the cerebral cortex and the cerebellum are best predicted by the semantic feature space. Therefore, voxels in these regions can be reliably interpreted to determine changes in semantic tuning across attentional states.

To determine how attention warps semantic representations, we first plotted the visual-semantic model weights in the cerebellum and cerebral cortex across the two attention conditions. Because the semantic feature space spans 985 dimensions, it is impractical to visualize the weight vector for each voxel. Thus, for visualization purposes, the dimensionality of the model weights was reduced. The model weights were projected onto the first three principal components of a group semantic space used in several previous studies (Huth et al., 2016; de Heer et al., 2017; LeBel et al., 2021; Deniz et al., 2023), and then each of the three principal components was assigned to the red, green, or blue color channel, respectively. Under this visualization procedure each voxel is associated with an RGB value, and the resulting color can be interpreted to indicate the semantic tuning of individual voxels. (Note that because the projection onto the first three principal components maps the model weights onto continuous color values, the resulting RGB colormap is continuous and not discrete.)

**Figure 4** shows the visual-semantic maps for the “attend humans” condition and the “attend vehicles” condition, plotted on the cerebral and cerebellar surfaces of one participant (see **Supplementary Figure S.3** and **Supplementary Figure S.4** for the other participants). The figure highlights several voxels with distinct patterns of semantic modulation across the two attention conditions. For example, the highlighted cerebral voxel near the supplementary eye field (SEF) shows a strong tuning shift towards the attended target. When the participant attends to humans, this voxel represents concepts such as “girl”, “guy”, and “kid”. When the participant attends to vehicles, the same voxel represents concepts such as “parking”and “truck”. Therefore, the semantic tuning of this voxel shifts towards the attended semantic concept. By contrast, the highlighted cerebellar voxel in eye motor regions (AEA) shows a tuning shift away from the attended semantic concept. When the participant attends to humans, this cerebellar voxel represents concepts such as “road” and “miles.” When the participant attends to vehicles, the same cerebellar voxel represents concepts such as “quest” and “fantasy.” These results suggest that attention warps semantic representations in both the cerebral cortex and the cerebellum, and that these changes can be interpreted in detail at the level of individual voxels.

**Figure 4.**
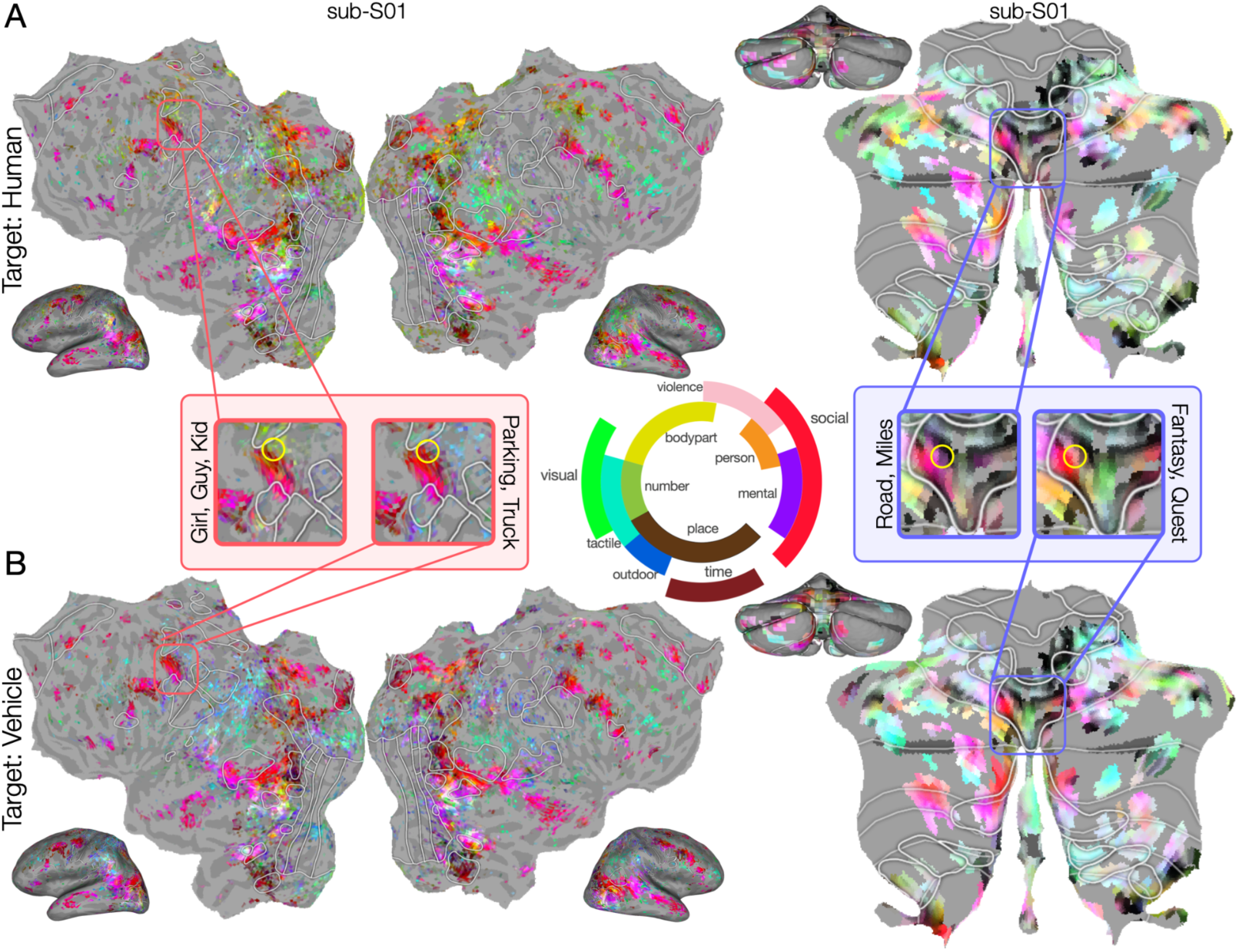
Cerebral and cerebellar visual-semantic maps recovered during the two attention conditions for one representative participant. To determine how attention warps semantic representations in the cerebral cortex and cerebellum, separate semantic maps were created for each attention condition. Because the semantic model weights are 985-dimensional, the model weight vector for each voxel was first projected onto a reduced three dimensional semantic space used in prior studies (Huth et al., 2016). An RGB colormap was then used to color each voxel according to its projection in this reduced semantic space. The color wheel in the center shows the major semantic categories associated with each color. **A.** Cortical and cerebellar semantic maps recovered during the “attend humans” condition for one representative participant. **B.** Cortical and cerebellar semantic maps recovered during the “attend vehicles” condition for the same participant. (**Supplementary Figure S.3** and **S.4** shows maps for the other participants.) The red inset shows a voxel in the cortex whose tuning shifts toward the attended semantic category. In the “attend humans” condition, this voxel represents concepts such as “girl” and “guy”. In the “attend vehicles” condition, the same voxel represents concepts such as “parking” and “truck”. The blue inset shows a voxel in the cerebellum whose tuning shifts away from the attended semantic category. In the “attend humans” condition, this voxel represents concepts such as “road” and “miles”. In the “attend vehicles” condition, the same voxel represents concepts such as “quest” and “fantasy”. These results suggest that attention warps semantic representations in both the cortex and the cerebellum, and that these changes can be interpreted in detail at the level of individual voxels.

To quantify the attentional tuning shift for each voxel in the cerebellum and cerebral cortex, we used a summary statistic called the Tuning Shift Index (TSI) (Çukur et al., 2013). The TSI quantifies both the magnitude and direction of tuning changes for each voxel. TSI values range from −1 to 1, where positive values correspond to tuning shifts towards the target, and negative values correspond to tuning shifts away from the target. **Figure 5A** and **5B** display the TSI values in the cerebral cortex and the cerebellar cortex for the group and for each individual participant. These maps reveal that cerebral voxels located in the intraparietal sulcus and regions of the frontal lobe shift towards the attended target, and cortical voxels located in most prefrontal regions and around the temporal parietal junction shift away from the attended target. Cerebellar voxels in lobules VI, VIIB, and along the vermis shift towards the attended target, and cerebellar voxels in medial parts of crus I/II and in lobules VIIIB and X shift away from the attended target. These results indicate that the attentional shifts are spatially heterogenous, and are distributed across many cortical and cerebellar regions that represent semantic information.

**Figure 5.**
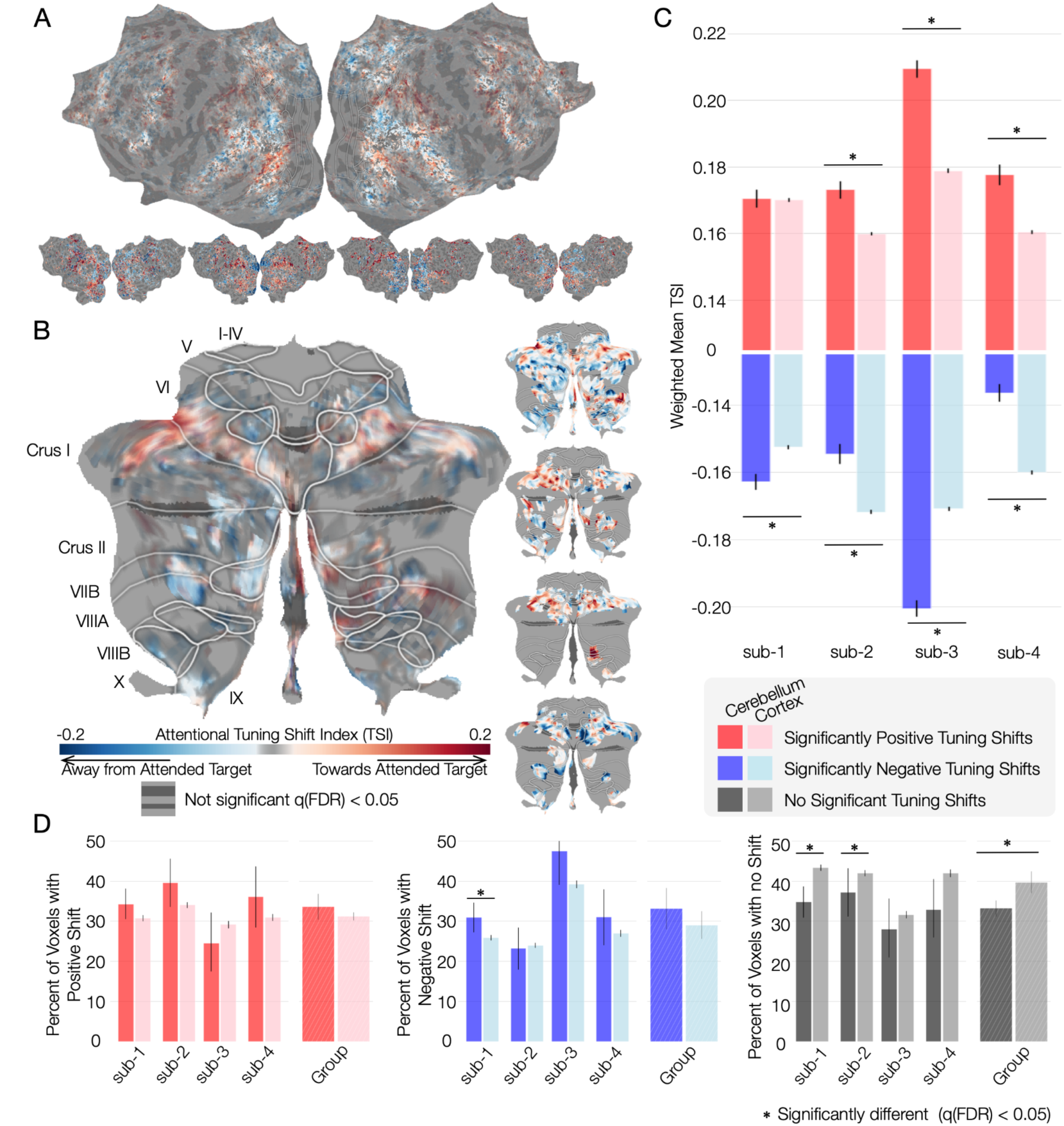
Semantic tuning shifts are larger and more widespread in the cerebellum than in the cerebral cortex. The previous results show qualitatively that attention shifts semantic tuning in both the cerebral cortex and the cerebellum. Here, we quantify these shifts to address the central question of how their magnitude and extent in the cerebellum compare to those in the cerebral cortex. To do so, the Tuning Shift Index (TSI) was calculated for each voxel (see Methods). **A.** Group-averaged and individual participant TSI maps in the cerebral cortex. **B.** Group-averaged and individual participant TSI maps in the cerebellum. Red indicates voxels that shift their semantic representations towards the attended target. Blue indicates voxels that shift their semantic representations away from the attended target. Gray indicates voxels that are not significantly predicted by the models in both conditions. Solid white lines indicate functionally localized ROIs. Group maps are thresholded to include only voxels with a significant tuning shift (q(FDR) < 0.05). Individual participant maps are thresholded to include only voxels with both significant model prediction accuracy and significant tuning shifts (both q(FDR) < 0.05). These maps reveal that both the cerebral cortex and the cerebellum show significant tuning shifts towards and away from the target in all participants. **C.** To directly compare the magnitude of tuning shifts in the cerebral cortex and the cerebellum, the average TSI was computed separately for voxels with positive (red) and negative (blue) tuning shifts. Only voxels with significant TSI values were included, and each voxel’s TSI was weighted by its semantic model prediction accuracy before averaging so that poorly predicted voxels would not bias the result. Bars show the weighted mean TSI for each participant, with lighter bars for the cerebral cortex and darker bars for the cerebellum. Error bars indicate SEM across voxels. Asterisks mark significant cortex–cerebellum differences (two-sided permutation test, q(FDR) < 0.05). Positive tuning shifts were significantly larger in the cerebellum than in the cortex for three of four participants. Negative tuning shifts were significantly larger in the cerebellum for two participants and in the cortex for the other two. These findings suggest that the magnitude of attentional shifts is larger in the cerebellum than in the cortex within areas that shift towards the attended target. **D.** To compare the extent of tuning shifts in the cerebral cortex and the cerebellum, the percentage of voxels showing a positive tuning shift (red), a negative tuning shift (blue), or no significant tuning shift (gray) was computed for each participant. Only voxels with significant model prediction accuracy (q(FDR) < 0.05) were included. Bars show these percentages for each participant and for the group, with lighter bars for the cerebral cortex and darker bars for the cerebellum. Error bars indicate SEM across voxels for individual participants and SEM across participant means for the group. Asterisks mark significant cerebellum-cortex differences (permutation test, q(FDR) < 0.05). Two participants had significantly more voxels with shifts towards the target in the cerebellum than in the cortex, and one participant had significantly more voxels with shifts away from the target in the cerebellum. Three of four participants had significantly fewer voxels with no tuning shift in the cerebellum than in the cortex, and this difference was also significant at the group level. This suggests that a larger proportion of voxels shift during attention in the cerebellum than in the cortex, with a bias towards shifts towards the target.

**Figure 5C** quantifies differences in attentional shifts between the two regions, showing the average positive TSI and the average negative TSI for each participant and each region. To ensure that poorly predicted voxels did not bias these values, the TSI value in each voxel was weighted by the semantic model prediction accuracy of that voxel prior to averaging, and only voxels with significant TSI values were considered (see **Methods**). This analysis reveals that the positive tuning shifts are consistently larger in the cerebellum than in the cerebral cortex (significant in three of the four participants, permutation test, q(FDR) < 0.05. The same trend is found in the fourth participant, but did not reach statistical significance.) By contrast, the pattern of negative tuning shifts was less consistent; two participants show significantly larger shifts in the cerebellum than in the cerebral cortex and two show significantly smaller shifts in the cerebellum than in the cerebral cortex (q(FDR) < 0.05). In summary, the magnitude of attentional shifts is consistently larger in the cerebellum than in the cerebral cortex only within areas that shift towards the attended target.

The finding that attentional shifts are larger in the cerebellum than in cerebral cortex may indicate that semantic representations in the cerebellum are affected by attention more strongly than are representations in the cortex. If this were the case, one would also expect to find a greater portion of the cerebellum to be modulated by attention compared to the cerebral cortex. **Figure 5D** shows the fraction of significantly predicted voxels (q(FDR) <0.05) that have positive or negative tuning shifts in both the cerebral cortex and the cerebellar cortex. The fraction of voxels with significant tuning shifts is significantly smaller in the cerebral cortex than in the cerebellar cortex. This effect is seen in all four individuals and is significant at the group level (q(FDR) < 0.05). However, when we perform this analysis separately for positive and negative voxels, there is no significant difference at the group level (p > 0.05). The consistent significant increase in magnitude of the tuning shifts at the individual level suggests that there are stronger attention effects in the cerebellum than in the cortex, particularly for positive tuning shifts.

## Discussion

Prior studies have shown that visual-semantic representations evoked by watching movies are distributed across both the cerebral cortex and the cerebellum (Huth et al., 2012, 2016; LeBel et al., 2021; Popham et al., 2021). These semantic representations in the cortex can be modulated by attentional state; some voxels shift their tuning toward the attended target, while others shift away from the attended target (Çukur et al., 2013; Shahdloo et al., 2020). Here we sought to determine how attention modulates the representation of semantic information in the cerebellum, and whether the direction and/or magnitude of the shifts differs from that observed in the cortex. To this end, we recorded brain activity while participants watched silent movie clips and selectively attended to either humans or vehicles. We fit voxelwise encoding models to map the representation of visual-semantic information under each attention condition, and directly compare attention-dependent tuning changes in the cerebellum and cortex.

We find that visual-semantic features are broadly represented in both the cerebral cortex and the cerebellum, and that much of this information is represented in cortical and cerebellar regions that are distinct from those encoding low-level visual and motor features. This pattern is similar to what has been observed in other movie-watching studies that focused on the cerebral cortex (Nishimoto et al., 2011; Huth et al., 2016; LeBel et al., 2021; Popham et al., 2021). Within the cerebellum, visual-semantic features reliably predict the BOLD response in bilateral Crus I/II and lobules VI/VIIB, regions that have been associated with language and cognitive control (Moberget et al., 2014; King et al., 2019; LeBel et al., 2021; Casto et al., 2025).

Turning to the core question of this study, we find that attention changes visual-semantic tuning across both the cortex and the cerebellum. In both structures about 60% of the voxels that were tuned for visual-semantics exhibited a significant shift, with about equal portions of voxels shifting tuning towards or away from the attended category. Although attentional tuning shifts have been reported previously in the cortex (Çukur et al., 2013; Shahdloo et al., 2020), to our knowledge this is the first time that such shifts have been observed in the cerebellum. This shows that cerebellar representations are a target of attentional control, and adds to work suggesting the cerebellum may also be a component of distributed cognitive control networks. Task-based studies consistently show cerebellar activity is modulated by attentional manipulations (Allen et al., 1997; Brissenden et al., 2021; Skipper and Lametti, 2021; Flierman et al., 2024; Qiu et al., 2026) and resting state fMRI studies identify cerebellar regions that correspond to the dorsal and ventral cortical resting states networks (Buckner et al., 2011; Marek et al., 2018). These cerebellar resting state attention networks are primarily in Lobules VI, VIIB, and VIIIA. Cerebellar voxels showing attentional modulation were observed in these regions, but also extended into Crus I and Crus II.

Prior neuropsychological studies have provided evidence that the cerebellum may contribute to top-down control. Deficits in cognitive control and visuospatial cognition are components of the cerebellar cognitive affective syndrome (CCAS), a condition observed in patients with focal or degenerative disorders of the cerebellum (Schmahmann and Sherman, 1998; Strick et al., 2009). For example, deficits are observed in studies examining task switching and attentional orienting studies (Akshoomoff and Courchesne, 1994; Townsend et al., 1999; Ravizza and Ivry, 2001; Bischoff-Grethe et al., 2002; Breska and Ivry, 2020, 2021), with the former associated with lesions in Crus I/II (Chirino-Pérez et al., 2021).The convergence between the neuroimaging and neuropsychological research raises the possibility that the cerebellum is causally involved in top-down cognitive control and attention, rather than only being subject to it.

This inference appears to conflict with prior studies that examined the effects of attention on cerebellar-dependent processes in sensorimotor control. That body of work has highlighted that the cerebellum can be insensitive to top-down information such as motor goals (Wolpert et al., 1998; Manto et al., 2012; Wang et al., 2024); for example, cerebellar-dependent adaptation continues to operate during motor tasks even when such adaptation makes performance worse (Mazzoni and Krakauer, 2006; Taylor et al., 2010; Taylor and Ivry, 2011). In contrast, our results show that an instructed attentional state can modulate the tuning of cerebellar voxels in a non-motor domain (Sokolov et al., 2017). To resolve this apparent conflict, future work should directly compare attentional modulation of cerebellar representations across motor and non-motor domains.

While the overall pattern of attentional modulation of visual-semantic representations was similar for the cortex and cerebellum, we also found some quantitative differences between these structures. First, a larger proportion of cerebellar voxels showed attentional modulation of visual-semantic representations than was found in the cerebral cortex. Second, for voxels in which visual-semantic tuning shifted towards the target, the magnitude of the tuning shifts tended to be larger in the cerebellum. The picture was more variable for voxels in which tuning shifted away from the target.

The precise reason why the modulatory effects of attention in the cerebellum are relatively larger than they are in the cortex is unclear. This effect might reflect intracerebellar computations that are modulated directly by attentional state. Alternatively, the cerebellar BOLD response might partially mirror attentional modulation of cortical regions that project to the cerebellum (Kelly and Strick, 1999; Shahshahani et al., 2024), or there could be some sort of multiplicative gain operating within the cerebellum that amplifies cortical signals depending on attentional state. Another possibility is that there are non-cortical sources of cerebellar input that are modulated by attention.

Finally, we note that the differences in the magnitude of attentional shifts between the cortex and the cerebellum might partly reflect unique methodological challenges associated with cerebellar fMRI, such as susceptibility related inhomogeneities and physiological noise properties that differ between the cerebellum and cortex (Diedrichsen et al., 2010; van der Zwaag et al., 2015). These methodological issues might have contributed to the differences between how attention modulated semantic representations in the cortex and cerebellum. In sum, adjudicating between these alternatives will require new data and computational models that can better explain which aspects of attentional modulation in the cerebellum are inherited from the cortex, and which aspects reflect local effects within the cerebellum.

What might be the advantage of large attentional shifts in the cerebellum? As noted in the Introduction, tuning shifts toward an attended target increase the effective resolution of the semantic representation, which might increase discriminability of the attended target from distractors. This would provide an efficient mechanism for rapidly amplifying goal-relevant features. Various lines of evidence highlight the importance of the cerebellum in generating predictions in motor, perceptual, and cognitive tasks (Perrett et al., 1993; O’Reilly et al., 2008; Breska and Ivry, 2020, 2021). Moreover, the predictive capability of the cerebellum is hypothesized to operate on a faster time scale than the cortex (Ivry, 1996; Sokolov et al., 2017; Boven et al., 2023). Therefore, the increase in attentional modulation in the cerebellum may improve the efficiency of semantic processing.

In summary, the present findings extend prior work by demonstrating that semantic representations within the cerebellum are modulated by attentional state. These goal-dependent tuning shifts provide novel evidence that cerebellar semantic representations are shaped by cognitive control, and suggest the cerebellum may contribute to that control.

## Supplementary Figures

**Figure S1.**
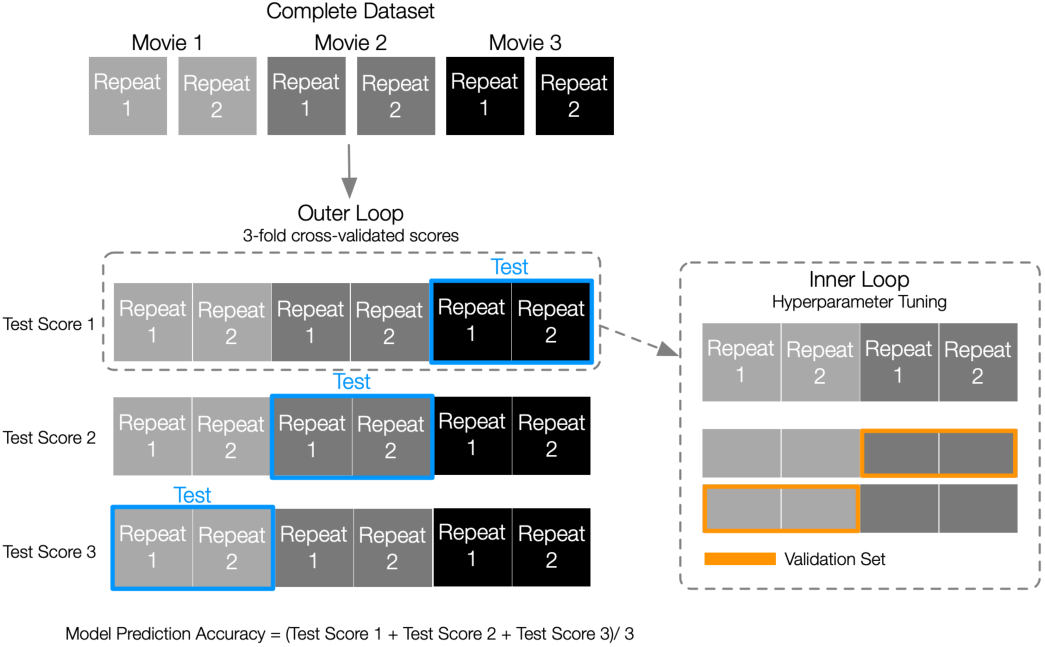
Nested Cross-validation scheme. Voxelwise encoding models were trained and evaluated using nested cross-validation. In each outer fold, data were split into training and test sets. Within the training set, inner folds were used to select voxel-specific regularization hyperparameters via random search (400 iterations across 20 log-spaced values between 10 and 10^20^). Models were refit on the outer training set using the best hyperparameters and evaluated on the held-out test set. Prediction accuracies (R²) were averaged across outer folds.

**Figure S2.**
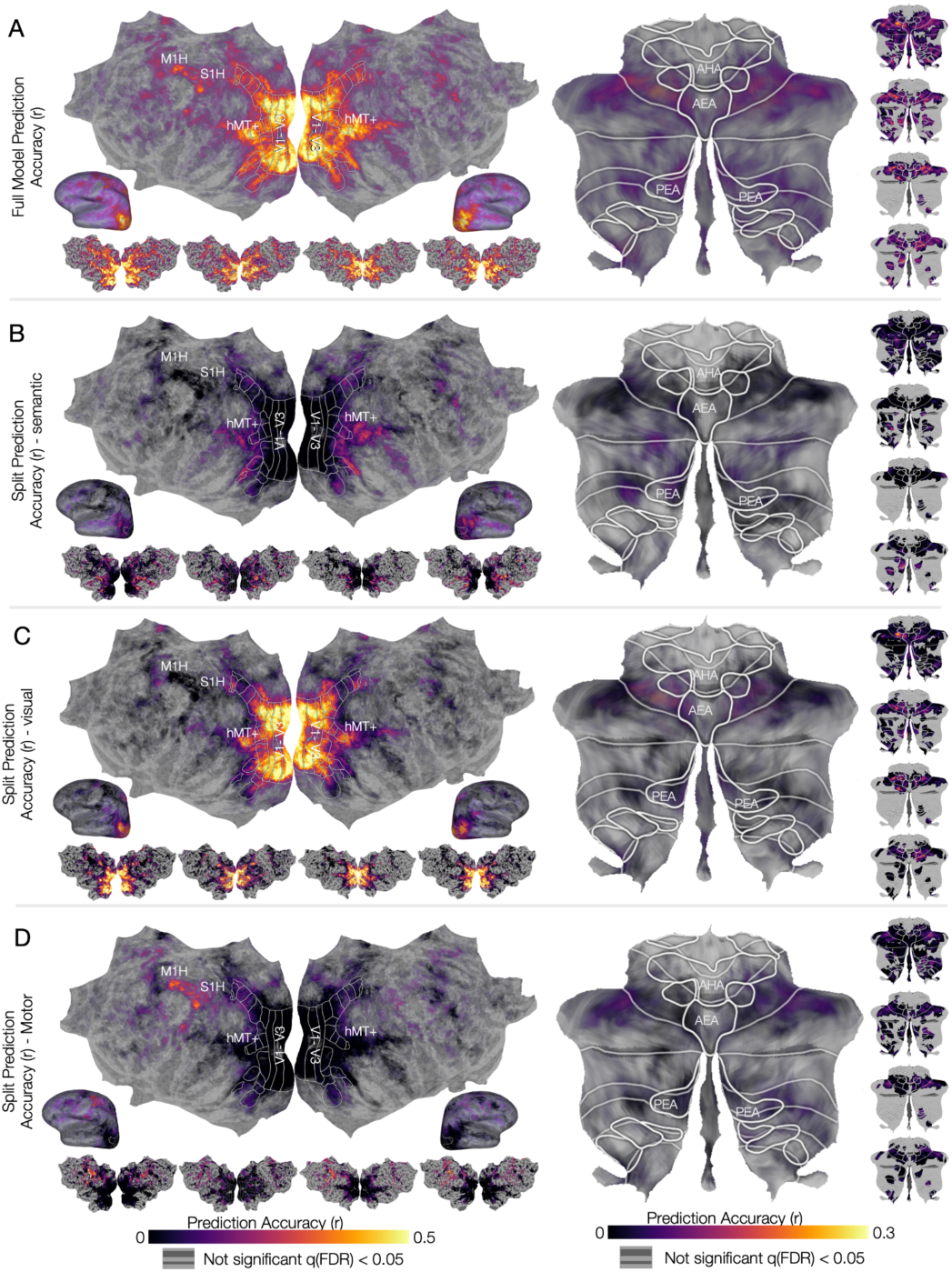
Prediction accuracy of voxelwise encoding models in the cerebral cortex and the cerebellar cortex in the “attend vehicle” condition. To identify regions of the cerebellum and cortex that selectively represent semantic information, banded ridge regression was used to fit voxelwise encoding models with three feature spaces: visual-semantic, low-level visual, and motor. A nested cross-validation procedure was used to train and test models in separate data sets. To identify cortical and cerebellar areas that are well predicted by the voxelwise encoding model, model prediction accuracy (correlation, *r*) was averaged across participants, and projected onto the fsaverage and SUIT surfaces respectively. A. Model prediction accuracy (correlation, *r*) for the full model, comprising all three feature spaces, is plotted on flattened cortical and cerebellar surfaces for the group (center, large) and in each individual participant (small). The full model in the cerebrum shows high prediction accuracy through the visual cortex, along the temporal lobe, and primary somatomotor cortex. Within the cerebellum, the full model shows high prediction accuracy in lobules VI, Crus I/II, and lobule VIIB. To disentangle the contribution of each feature space to model predictions, the split prediction accuracy was plotted on the same cortical and cerebellar surfaces (B, C, and D). The split prediction accuracy can be interpreted as the variance explained by each feature space. **B.** Split prediction accuracy maps for the visual-semantic feature space. The visual-semantic features best explain variance in the temporal and frontal lobes in cerebral cortex, and in Crus I/II and lobule VIIB in the cerebellum. **C.** Split prediction accuracy maps for the low-level visual feature space. The low-level visual feature space best explains variance in the visual cortex, and in lobules VI and vermis in the cerebellum. **D.** Split prediction accuracy maps for the motor feature space. The motor feature space best explains variance in the left primary somatomotor cortex, and in right lobules I-IV and VI in the cerebellum. These results indicate that separate regions in the cortex and cerebellum represent semantic, low-level visual, and motor information during the naturalistic visual search task. Representations in voxels that are well predicted by the visual-semantic feature space can be reliably interpreted to determine changes in semantic tuning across attentional states.

**Figure S3.**
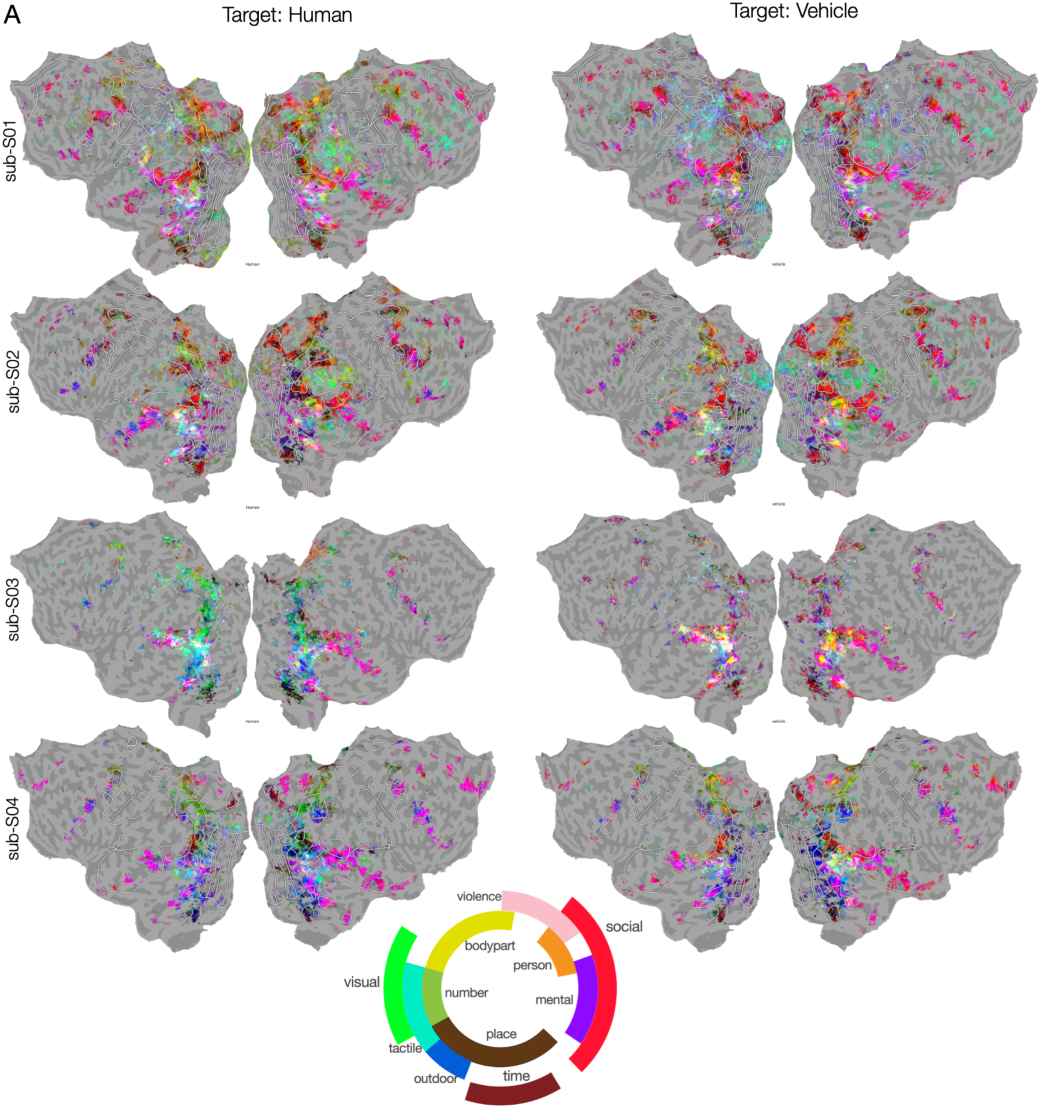
Cerebral visual-semantic maps recovered during the two attention conditions for all participants. To determine how attention warps semantic representations in the cerebral cortex, separate semantic maps were created for each attention condition. Because the semantic model weights are 985-dimensional, the model weight vector for each voxel was first projected onto a reduced three dimensional semantic space used in prior studies (Huth et al., 2016). An RGB colormap was then used to color each voxel according to its projection in this reduced semantic space. The color wheel in the center shows the major semantic categories associated with each color. **A.** Cerebral semantic maps recovered during the “attend humans” and “attend vehicles” condition for all participants. These results suggest that attention warps semantic representations in the cortex, and that these changes can be interpreted in detail at the level of individual voxels.

**Figure S4.**
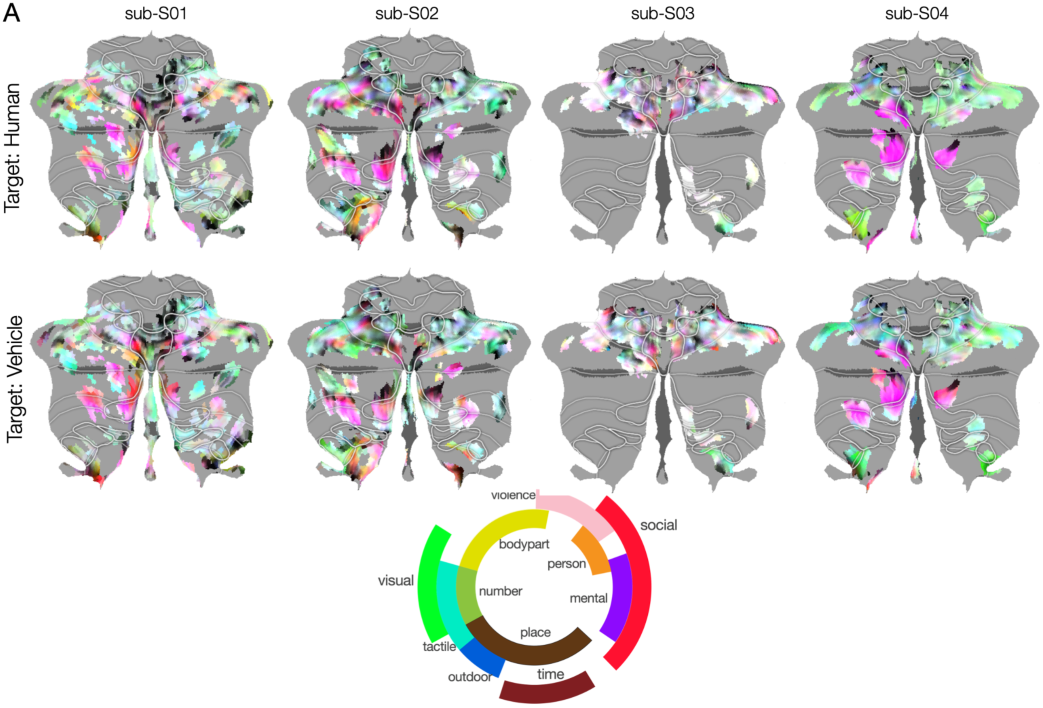
Cerebellar visual-semantic maps recovered during the two attention conditions for all participants. To determine how attention warps semantic representations in the cerebellar cortex, separate semantic maps were created for each attention condition. Because the semantic model weights are 985-dimensional, the model weight vector for each voxel was first projected onto a reduced three dimensional semantic space used in prior studies (Huth et al., 2016). An RGB colormap was then used to color each voxel according to its projection in this reduced semantic space. The color wheel in the center shows the major semantic categories associated with each color. **A.** Cerebellar semantic maps recovered during the “attend humans” and “attend vehicles” condition for all participants. These results suggest that attention warps semantic representations in the cerebellum, and that these changes can be interpreted in detail at the level of individual voxels.

**Figure S5.**
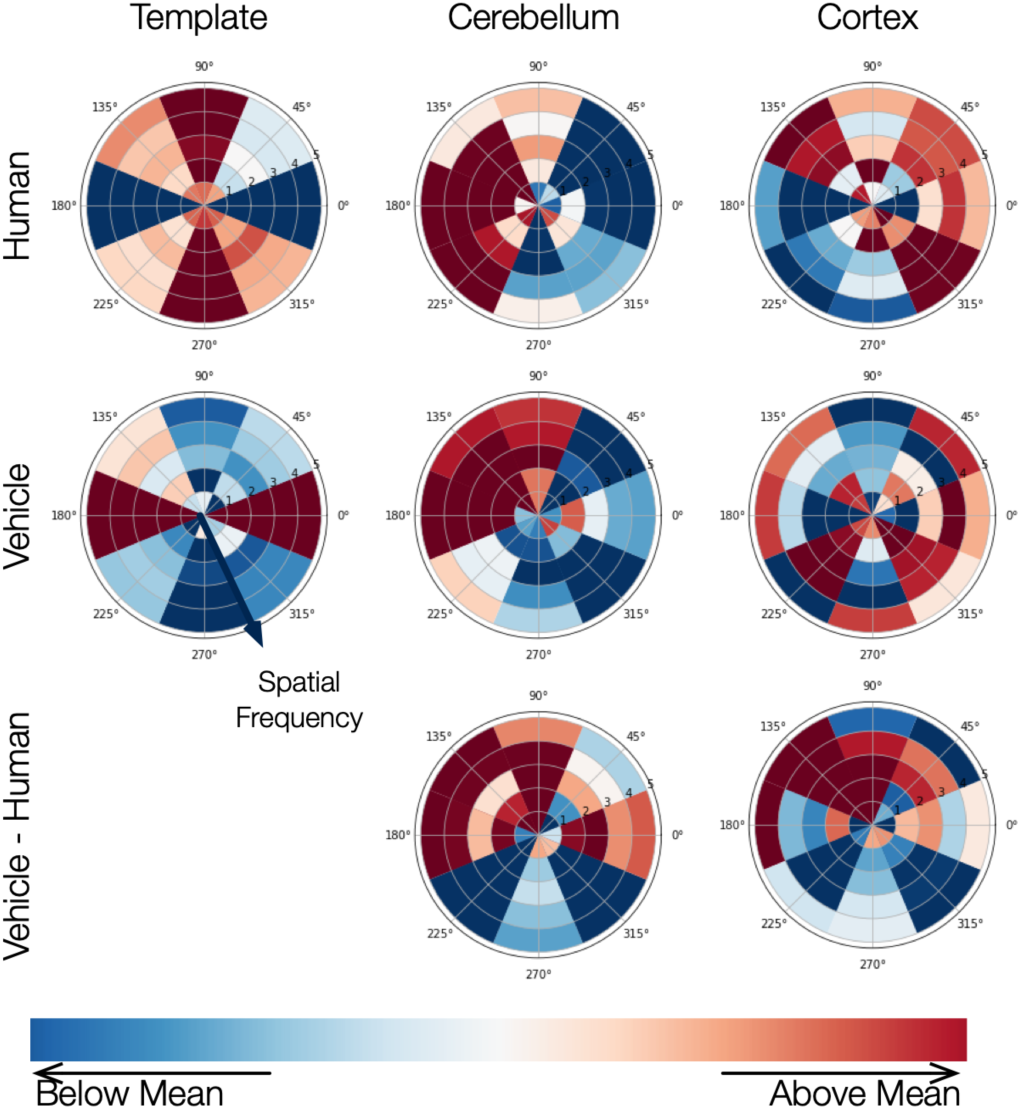
Motion-energy tuning shifts in the cortex and the cerebellum. To test for potential tuning shifts of motion energy representations in all voxels across participants, we generated a template for low-level visual features across the two attentional targets. This was done by calculating the average motion-energy features for all frames that included the target. The templates for the targets show opposite patterns: the human template is biased toward vertical orientations across spatial frequencies. while the vehicle template is biased toward horizontal orientations across spatial frequencies. We then looked at the motion-energy tuning for all voxels in the cerebellum and cortex. No obvious patterns were seen or large changes. However this could be improved by doing the analysis voxelwise.

## Notes

This work was supported by the National Science Foundation (DGE 2146752), the National Institutes of Health (R35NS116883, R01DC0170941), and the Office of Naval Research (N00014-20-1-2002, N00014-22-1-2217).

### Competing Interest Statement

The authors have declared no competing interest.

## Works Cited

Adelson EH, Bergen JR (1985) Spatiotemporal energy models for the perception of motion. J Opt Soc Am A 2:284–299.

Akshoomoff NA, Courchesne E (1994) ERP evidence for a shifting attention deficit in patients with damage to the cerebellum. J Cogn Neurosci 6:388–399.

Allen G, Buxton RB, Wong EC, Courchesne E (1997) Attentional activation of the cerebellum independent of motor involvement. Science 275:1940–1943.

Andersson JLR, Skare S, Ashburner J (2003) How to correct susceptibility distortions in spin-echo echo-planar images: application to diffusion tensor imaging. Neuroimage 20:870–888.

Avants BB, Tustison NJ, Song G, Cook PA, Klein A, Gee JC (2011) A reproducible evaluation of ANTs similarity metric performance in brain image registration. Neuroimage 54:2033–2044.

Behzadi Y, Restom K, Liau J, Liu TT (2007) A component based noise correction method (CompCor) for BOLD and perfusion based fMRI. Neuroimage 37:90–101.

Benjamini Y, Hochberg Y (1995) Controlling the False Discovery Rate: A Practical and Powerful Approach to Multiple Testing. J R Stat Soc Series B Stat Methodol 57:289–300.

Bischoff-Grethe A, Ivry RB, Grafton ST (2002) Cerebellar involvement in response reassignment rather than attention. J Neurosci 22:546–553.

Boven E, Pemberton J, Chadderton P, Apps R, Costa RP (2023) Cerebro-cerebellar networks facilitate learning through feedback decoupling. Nat Commun 14:51.

Breska A, Ivry RB (2020) Context-specific control over the neural dynamics of temporal attention by the human cerebellum. Sci Adv 6 Available at: 10.1126/sciadv.abb1141.

Breska A, Ivry RB (2021) The human cerebellum is essential for modulating perceptual sensitivity based on temporal expectations. Elife 10 Available at: 10.7554/eLife.66743.

Brissenden JA, Tobyne SM, Halko MA, Somers DC (2021) Stimulus-specific visual working memory representations in human cerebellar lobule VIIb/VIIIa. J Neurosci 41:1033–1045.

Buckner RL, Krienen FM, Castellanos A, Diaz JC, Yeo BTT (2011) The organization of the human cerebellum estimated by intrinsic functional connectivity. J Neurophysiol 106:2322–2345.

Caligiore D, Pezzulo G, Baldassarre G, Bostan AC, Strick PL, Doya K, Helmich RC, Dirkx M, Houk J, Jörntell H, Lago-Rodriguez A, Galea JM, Miall RC, Popa T, Kishore A, Verschure PFMJ, Zucca R, Herreros I (2017) Consensus Paper: Towards a Systems-Level View of Cerebellar Function: the Interplay Between Cerebellum, Basal Ganglia, and Cortex. Cerebellum 16:203–229.

Casto C, Small H, Poliak M, Tuckute G, Lipkin B, Wolna A, D’Mello AM, Fedorenko E (2025) The cerebellar components of the human language network. bioRxivorg:2025.04.14.645351 Available at: https://www.biorxiv.org/content/10.1101/2025.04.14.645351v1.abstract [Accessed April 21, 2025].

Chirino-Pérez A, Marrufo-Meléndez OR, Muñoz-López JI, Hernandez-Castillo CR, Ramirez-Garcia G, Díaz R, Nuñez-Orozco L, Fernandez-Ruiz J (2021) Mapping the Cerebellar Cognitive Affective Syndrome in Patients with Chronic Cerebellar Strokes. Cerebellum Available at: 10.1007/s12311-021-01290-3.

Çukur T, Nishimoto S, Huth AG, Gallant JL (2013) Attention during natural vision warps semantic representation across the human brain. Nat Neurosci 16:763–770.

Dale AM, Fischl B, Sereno MI (1999) Cortical surface-based analysis. I. Segmentation and surface reconstruction. Neuroimage 9:179–194.

David SV, Hayden BY, Mazer JA, Gallant JL (2008) Attention to stimulus features shifts spectral tuning of V4 neurons during natural vision. Neuron 59:509–521.

de Heer WA, Huth AG, Griffiths TL, Gallant JL, Theunissen FE (2017) The Hierarchical Cortical Organization of Human Speech Processing. J Neurosci 37:6539–6557.

Deniz F, Tseng C, Wehbe L, Dupré la Tour T, Gallant JL (2023) Semantic representations during language comprehension are affected by context. J Neurosci Available at: 10.1523/JNEUROSCI.2459-21.2023.

Diedrichsen J (2006) A spatially unbiased atlas template of the human cerebellum. Available at: 10.1016/j.neuroimage.2006.05.056.

Diedrichsen J, Verstynen T, Schlerf J, Wiestler T (2010) Advances in functional imaging of the human cerebellum. Curr Opin Neurol 23:382–387.

D’Mello AM, Centanni TM, Gabrieli JDE, Christodoulou JA (2020) Cerebellar contributions to rapid semantic processing in reading. Brain Lang 208:104828.

Downing PE, Jiang Y, Shuman M, Kanwisher N (2001) A cortical area selective for visual processing of the human body. Science 293:2470–2473.

Dupre la Tour T, Eickenberg M, Gallant JL (2022) Feature-space selection with banded ridge regression. bioRxiv:2022.05.05.490831 Available at: https://www.biorxiv.org/content/10.1101/2022.05.05.490831v1 [Accessed May 6, 2022].

Dupré la Tour T, Eickenberg M, Nunez-Elizalde AO, Gallant JL (2022) Feature-space selection with banded ridge regression. Neuroimage 264:119728.

Dupré la Tour T, Visconti di Oleggio Castello M, Gallant JL (2025) The Voxelwise Encoding Model framework: a tutorial introduction to fitting encoding models to fMRI data. Available at: https://direct.mit.edu/imag/article/doi/10.1162/imag_a_00575/128936.

Epstein R, Kanwisher N (1998) A cortical representation of the local visual environment. Nature 392:598–601.

Fellbaum C, Miller GA (2021) WordNet. MIT Press Available at: https://mitpress.mit.edu/9780262561167/ [Accessed November 28, 2023].

Flierman NA, Koay SA, van Hoogstraten WS, Ruigrok TJH, Roelfsema PR, Badura A, De Zeeuw CI (2024) Encoding of cerebellar dentate neuronal activity during visual attention in rhesus macaques. bioRxiv:2024.07.18.604119 Available at: https://www.biorxiv.org/content/10.1101/2024.07.18.604119v1.abstract [Accessed July 25, 2024].

Gao JS, Huth AG, Lescroart MD, Gallant JL (2015) Pycortex: an interactive surface visualizer for fMRI. Front Neuroinform 9:23.

Gorgolewski K, Burns CD, Madison C, Clark D, Halchenko YO, Waskom ML, Ghosh SS (2011) Nipype: a flexible, lightweight and extensible neuroimaging data processing framework in python. Front Neuroinform 5:13.

Gorgolewski KJ et al. (2016) Nipype: a flexible, lightweight and extensible neuroimaging data processing framework in Python. 0.12.0-rc1. Available at: 10.5281/zenodo.50186.

Greve DN, Fischl B (2009) Accurate and robust brain image alignment using boundary-based registration. Neuroimage 48:63–72.

Han S, Carass A, He Y, Prince JL (2020) Automatic cerebellum anatomical parcellation using U-Net with locally constrained optimization. Neuroimage 218:116819.

Hansen KA, Kay KN, Gallant JL (2007) Topographic organization in and near human visual area V4. J Neurosci 27:11896–11911.

Huth AG, de Heer WA, Griffiths TL, Theunissen FE, Gallant JL (2016) Natural speech reveals the semantic maps that tile human cerebral cortex. Nature 532:453–458.

Huth AG, Nishimoto S, Vu AT, Gallant JL (2012) A continuous semantic space describes the representation of thousands of object and action categories across the human brain. Neuron 76:1210–1224.

Ignashchenkova A, Dicke PW, Haarmeier T, Thier P (2004) Neuron-specific contribution of the superior colliculus to overt and covert shifts of attention. Nat Neurosci 7:56–64.

Ivry RB (1996) The representation of temporal information in perception and motor control. Curr Opin Neurobiol 6:851–857.

Jenkinson M, Bannister P, Brady M, Smith S (2002) Improved Optimization for the Robust and Accurate Linear Registration and Motion Correction of Brain Images. Neuroimage 17:825–841.

Kanwisher N, McDermott J, Chun MM (1997) The fusiform face area: a module in human extrastriate cortex specialized for face perception. J Neurosci 17:4302–4311.

Kastner S, De Weerd P, Desimone R, Ungerleider LG (1998) Mechanisms of directed attention in the human extrastriate cortex as revealed by functional MRI. Science 282:108–111.

Kelly RM, Strick PL (1999) Cerebellar Loops with Motor Cortex and Prefrontal Cortex of a Nonhuman Primate. Journal of Neuroscience 1994:8432–8444.

King M, Hernandez-Castillo CR, Poldrack RA, Ivry RB, Diedrichsen J (2019) Functional boundaries in the human cerebellum revealed by a multi-domain task battery. Nat Neurosci 22:1371–1378.

LeBel A, D’Mello AM (2023) A seat at the (language) table: incorporating the cerebellum into frameworks for language processing. Current Opinion in Behavioral Sciences 53:101310.

LeBel A, Jain S, Huth AG (2021) Voxelwise Encoding Models Show That Cerebellar Language Representations Are Highly Conceptual. J Neurosci 41:10341–10355.

LeBel A, Wagner L, Jain S, Adhikari-Desai A, Gupta B, Morgenthal A, Tang J, Xu L, Huth AG (2023) A natural language fMRI dataset for voxelwise encoding models. Scientific Data 10:1–12.

Le TH, Pardo JV, Hu X (1998) 4 T-fMRI study of nonspatial shifting of selective attention: cerebellar and parietal contributions. J Neurophysiol 79:1535–1548.

Manto M et al. (2012) Consensus paper: roles of the cerebellum in motor control--the diversity of ideas on cerebellar involvement in movement. Cerebellum 11:457–487.

Marek S et al. (2018) Spatial and Temporal Organization of the Individual Human Cerebellum. Neuron 100:977–993.e7.

Mazer JA, Gallant JL (2003) Goal-related activity in V4 during free viewing visual search. Evidence for a ventral stream visual salience map. Neuron 40:1241–1250.

Mazzoni P, Krakauer JW (2006) An implicit plan overrides an explicit strategy during visuomotor adaptation. J Neurosci 26:3642–3645.

Meschke EX, Visconti di Oleggio Castello M, Dupré la Tour T, Gallant JL (2023) Model connectivity: leveraging the power of encoding models to overcome the limitations of functional connectivity. bioRxiv:2023.07.17.549356 Available at: https://www.biorxiv.org/content/10.1101/2023.07.17.549356v1.abstract [Accessed October 17, 2023].

Moberget T, Gullesen EH, Andersson S, Ivry RB, Endestad T (2014) Generalized role for the cerebellum in encoding internal models: evidence from semantic processing. Journal of Neuroscience 34:2871–2878.

Murdoch BE (2010) The cerebellum and language: Historical perspective and review. Cortex 46:858–868.

Naselaris T, Kay KN, Nishimoto S, Gallant JL (2011) Encoding and decoding in fMRI. Neuroimage 56:400–410.

Nishimoto S, Vu AT, Naselaris T, Benjamini Y, Yu B, Gallant JL (2011) Reconstructing visual experiences from brain activity evoked by natural movies. Curr Biol 21:1641–1646.

Nunez-Elizalde AO, Huth AG, Gallant JL (2019) Voxelwise encoding models with non-spherical multivariate normal priors. Neuroimage 197:482–492.

O’Reilly JX, Mesulam MM, Nobre AC (2008) The cerebellum predicts the timing of perceptual events. J Neurosci 28:2252–2260.

Perrett SP, Ruiz BP, Mauk MD (1993) Cerebellar cortex lesions disrupt learning-dependent timing of conditioned eyelid responses. J Neurosci 13:1708–1718.

Popham SF, Huth AG, Bilenko NY, Deniz F, Gao JS, Nunez-Elizalde AO, Gallant JL (2021) Visual and linguistic semantic representations are aligned at the border of human visual cortex. Nat Neurosci 24:1628–1636.

Qiu BS, Woodard KM, Leonard B, Murray SO (2026) Stimulus-independent attentional modulation in cerebellar visual regions. J Neurosci:e0670262026.

Ravizza SM, Ivry RB (2001) Comparison of the basal ganglia and cerebellum in shifting attention. J Cogn Neurosci 13:285–297.

Saalmann YB, Pinsk MA, Wang L, Li X, Kastner S (2012) The pulvinar regulates information transmission between cortical areas based on attention demands. Science 337:753–756.

Schmahmann JD (2019) The cerebellum and cognition. Neurosci Lett 688:62–75.

Schmahmann JD, Sherman JC (1998) The cerebellar cognitive affective syndrome. Brain 121:561–579.

Shahdloo M, Çelik E, Çukur T (2020) Biased competition in semantic representation during natural visual search. Neuroimage 216:116383.

Shahshahani L, King M, Nettekoven C, Ivry RB, Diedrichsen J (2024) Selective recruitment of the cerebellum evidenced by task-dependent gating of inputs. Elife 13 Available at: https://elifesciences.org/articles/96386 [Accessed November 8, 2024].

Skipper JI, Lametti DR (2021) Speech Perception under the Tent: A Domain-general Predictive Role for the Cerebellum. J Cogn Neurosci 33:1517–1534.

Sokolov AA, Miall RC, Ivry RB (2017) The Cerebellum: Adaptive Prediction for Movement and Cognition. Trends Cogn Sci 21:313–332.

Stoodley CJ, Schmahmann JD (2010) Evidence for topographic organization in the cerebellum of motor control versus cognitive and affective processing. Cortex 46:831–844.

Strick PL, Dum RP, Fiez JA (2009) Cerebellum and nonmotor function. Annu Rev Neurosci 32:413–434.

St-Yves G, Naselaris T (2018) The feature-weighted receptive field: an interpretable encoding model for complex feature spaces. Neuroimage 180:188–202.

Taylor JA, Ivry RB (2011) Flexible cognitive strategies during motor learning. PLoS Comput Biol 7:e1001096.

Taylor JA, Klemfuss NM, Ivry RB (2010) An explicit strategy prevails when the cerebellum fails to compute movement errors. Cerebellum 9:580–586.

Townsend J, Courchesne E, Covington J, Westerfield M, Harris NS, Lyden P, Lowry TP, Press GA (1999) Spatial attention deficits in patients with acquired or developmental cerebellar abnormality. J Neurosci 19:5632–5643.

van der Zwaag W, Jorge J, Butticaz D, Gruetter R (2015) Physiological noise in human cerebellar fMRI. MAGMA 28:485–492.

van Es DM, van der Zwaag W, Knapen T (2019) Topographic Maps of Visual Space in the Human Cerebellum. Curr Biol 0 Available at: 10.1016/j.cub.2019.04.012.

Visconti di Oleggio Castello M, Borghesani V, Rankin KP, Gallant JL, Vu AT (2021) Functional imaging of the anterior temporal lobe: effects of acceleration, multi-echo acquisition, and reconstruction methods. In: International Society for Magentic Resonance in Medicine.

Wang T, Avraham G, Tsay JS, Abram SJ, Ivry RB (2024) Perturbation variability does not influence implicit sensorimotor adaptation. PLoS Comput Biol 20:e1011951.

Watson AB, Ahumada AJ Jr (1985) Model of human visual-motion sensing. J Opt Soc Am A 2:322–341.

Wolpert DM, Miall RC, Kawato M (1998) Internal models in the cerebellum. Trends Cogn Sci 2:338–347.

Woolrich MW, Jbabdi S, Patenaude B, Chappell M, Makni S, Behrens T, Beckmann C, Jenkinson M, Smith SM (2009) Bayesian analysis of neuroimaging data in FSL. Neuroimage 45:S173–S186.

